# Synaptic density and neuronal metabolic function measured by PET in the unilateral 6-OHDA rat model of Parkinson’s disease

**DOI:** 10.1101/2021.05.27.444950

**Authors:** Nakul Ravi Raval, Frederik Gudmundsen, Morten Juhl, Ida Vang Andersen, Nikolaj Raahauge Speth, Annesofie Videbæk, Ida Nymann Petersen, Jens Damsgaard Mikkelsen, Patrick MacDonald Fisher, Matthias Manfred Herth, Pontus Plavén-Sigray, Gitte Moos Knudsen, Mikael Palner

**Author notes:** **Correspondence:** Mikael Palner, Neurobiology Research Unit, Section 8057, Rigshospitalet, Blegdamsvej 9, DK-2100 Copenhagen Ø, Denmark. These authors have contributed equally to this work and share the first authorship (Nakul Ravi Raval, Frederik Gudmundsen).

## Abstract

Parkinson’s disease (PD) is caused by progressive neurodegeneration and characterised by motor dysfunction. Neurodegeneration of dopaminergic neurons also causes aberrations within the cortico-striato-thalamo-cortical (CSTC) circuit, which has been hypothesised to lead to non-motor symptoms such as depression. Individuals with PD have both lower synaptic density and changes in neuronal metabolic function in the basal ganglia, as measured using [^11^C]UCB-J and [^18^F]FDG positron emission tomography (PET), respectively. However, the two radioligands have not been directly compared in the same PD subject or in neurodegeneration animal models. Here, we investigate [11C]UCB-J binding and [^18^F]FDG uptake in the CSTC circuit following a unilateral dopaminergic lesion in rats and compare it to sham lesioned rats.

Rats received either a unilateral injection of 6-hydroxydopamine (6-OHDA) or saline in the medial forebrain bundle and rostral substantia nigra (n=4/group). After three weeks, all rats underwent two PET scans using [^18^F]FDG, followed by [^11^C]UCB-J on a separate day. [^18^F]FDG uptake and [^11^C]UCB-J binding were both lower in the ipsilateral striatal regions compared to the contralateral regions. Using [^11^C]UCB-J, we could detect an 8.7% decrease in the ipsilateral ventral midbrain, compared to a 2.9% decrease in ventral midbrain using [^18^F]FDG. Differential changes between hemispheres for [^11^C]UCB-J and [^18^F]FDG outcomes were also evident in the CSTC circuit’s cortical regions, especially in the orbitofrontal cortex and medial prefrontal cortex where higher synaptic density yet lower neuronal metabolic function was observed, following lesioning.

In conclusion, [^11^C]UCB-J and [^18^F]FDG PET can detect divergent changes following a dopaminergic lesion in rats, especially in cortical regions that are not directly affected by the neurotoxin. These results suggest that combined [^11^C]UCB-J and [^18^F]FDG scans could yield a better picture of the heterogeneous cerebral changes in neurodegenerative disorders.

## 1 Introduction

Several techniques have been developed to identify disease-related neuronal patterns to aid early detection and differential diagnoses of Parkinson’s disease (PD). Examples of such methods are positron emission tomography (PET) imaging to measure glucose metabolism(Loane and Politis, 2011), dopamine synthesis, transporters, or receptors (Kerstens and Varrone, 2020). In PD, one affected neuronal circuit is the cortico-striato-thalamo-cortical (CSTC) circuit (Vriend et al., 2014). The CSTC circuit connects the cortex with the basal ganglia to control and coordinate goal-directed behaviour. This circuit can be further divided into three loops: the motor, limbic, and associative circuits (Groenewegen and Trimble, 2007; Vriend et al., 2014). The dopamine system innervates the striatal regions of the CSTC circuits and is critical in modulating their output. A model of 6-hydroxydopamine (6-OHDA) induced dopaminergic lesion leads to modulation within the CSTC, which will further help understand this circuit (Schwarting and Huston, 1996).

[^11^C]UCB-J is a PET radioligand showing high affinity to synaptic vesicle glycoprotein 2A (SV2A) (Nabulsi et al., 2016). SV2A is ubiquitously expressed throughout the brain (Bajjalieh et al., 1994; Südhof, 2004) and is a suitable proxy for synaptic density (Finnema et al., 2016). Accordingly, [^11^C]UCB-J PET may serve as a biomarker in neurodegenerative disorders, where the loss of synapses is thought to play a vital role in the pathophysiology (Holland et al., 2020; Matuskey et al., 2020; Mecca et al., 2020; Nicastro et al., 2020; Wilson et al., 2020; O’Dell et al., 2021). [^18^F]fluorodeoxyglucose (FDG) is a glucose analogue used to measure neuronal glucose consumption and metabolic function. [^18^F]FDG PET has also been used as a surrogate marker for neuronal integrity and function (Mosconi, 2013). Only very recently, [^18^F]FDG and [^11^C]UCB-J were tested in the same Alzheimer patients (Chen et al., 2021), where [^11^C]UCB-J proved valuable as a clinical tracer and marker for disease progression, which may be helpful in drug development. This combination of radioligands has not been tested in human PD subjects or animal models of neurodegeneration.

Here we present a multimodal PET study using dynamic [^11^C]UCB-J and static [^18^F]FDG scans in the rat model of 6-OHDA severe unilateral-dopaminergic lesioning induced by combined unilateral 6-OHDA injection in the medial forebrain bundle and rostral substantia nigra (Yuan et al., 2005; Blandini et al., 2008). We have previously shown that 6-OHDA lesioning lowers postsynaptic dopamine receptor density and presynaptic capacity to release amphetamine (Palner et al., 2011). Thus, we hypothesise that the loss of dopaminergic neurons will cause a decrease in [^11^C]UCB-J binding and [^18^F]FDG uptake, especially in the ipsilateral basal ganglia (substantia nigra, ventral tegmental area, whole striatum, dorsolateral striatum, dorsomedial striatum, and nucleus accumbens). Furthermore, we compare the effect sizes of [^18^F]FDG uptake and [^11^C]UCB-J binding to detect changes after a unilateral dopaminergic lesioning of the rat brain. As a control to assess differential changes, we used both the contralateral hemisphere and compared the 6-OHDA model to a group of sham-lesioned rats. Several studies have successfully detected changes in regional [^18^F]FDG uptake after a 6-OHDA lesion in both rats (Casteels et al., 2008; Jang et al., 2012; Silva et al., 2013; Kordys et al., 2017) and mice (Im et al., 2016). One recent study has also performed [^11^C]UCB-J PET in the 6-OHDA lesion model, although with some methodological differences (Thomsen et al., 2021b).

The results of our study indicate that dopaminergic lesions lead to a loss of presynaptic density in the striatal regions, as measured by [^11^C]UCB-J, which is similar to changes in neuronal metabolic function, as measured by [^18^F]FDG. Interestingly, the dopaminergic lesion caused divergent changes between the two radioligands in cortical regions of the CSTC circuit.

## 2 Materials and Methods

### 2.1 Animals

Eight female Long-Evans WT rats (239 ± 12 g, 10-11 weeks old when scanned) (Janvier) were used in this study. The animals were held under standard laboratory conditions with 12-hour light/12-hour dark cycles and ad libitum access to food and water. All animal experiments conformed to the European Commission’s Directive 2010/63/EU with approval from the Danish Council of Animal Ethics (Journal no. 2017-15-0201-01375) and the Department of Experimental Medicine, University of Copenhagen.

### 2.2 Stereotactic surgery and 6-OHDA lesion

The animals were acclimatised in the surgery room for at least 1 hour. Analgesia was provided with carprofen (Rimadyl, Zoetis, NJ, USA) 5 mg/kg, subcutaneous (SC), 45 minutes before the surgery and 24 hours and 48 hours postoperative. Before commencing the surgery, animals received desmethylimipramine (25 mg/kg, intraperitoneal (IP)) mixed in physiological saline. Desmethylimipramine protects the noradrenergic neurons from the neurotoxic effects (Esteban et al., 1999). Anaesthesia was induced with 3% isoflurane in oxygen and maintained through surgery with 1.2–1.8% isoflurane in oxygen. The rats were fixed on a stereotaxic apparatus (Kopf Instruments, Tujunga, CA, USA) with the incisor bar set 3.3 mm below the level of the ear bars. An incision was made on the scalp, and two bur-holes were drilled on one side of the skull using a dental micromotor and round bur (0.5 mm). A 2 µg/µL solution of 6-OHDA (2,5-Dihydroxytyramine hydrobromide, Sigma-Aldrich, Søborg, Denmark) in physiological saline containing 0.02% ascorbic acid or physiological saline (containing 0.02% ascorbic acid) was drawn into a 10 µL syringe with a 33 g needle (World Precision Instruments, Sarasota, FL, USA). 3 µL were infused into the medial forebrain bundle (coordinates: AP= 4.8 mm, ML= 1.7 mm, DV= 8 mm) and 3 µL infused rostral to substantia nigra (coordinates: AP= 3.6 mm, ML= 2 mm, DV= 8.3 mm) relative to the bregma to ensure unilateral dopaminergic degeneration. The infusion was delivered at 151 nL/minutes driven by an infusion pump (World Precision Instruments, Sarasota, FL, USA), followed by a 7 minute pause prior to a slow withdrawal of the syringe needle. The incision was sutured back. After recovery from anaesthesia, rats were returned to the recovery cage and housed alone for 48 hours and then housed in pairs for recovery of 21 days to allow the development of the lesions.

### 2.3 Study design and confirmation of lesion

Four rats were injected unilaterally with 6-OHDA, while another four were injected with physiological saline and divided into two groups, i.e., dopamine lesioned and sham lesioned; Figure 1 shows the study’s overall design. After the recovery period, the rats were subjected to two PET scans [^18^F]FDG at day 21 and [^11^C]UCB-J at approximately day 23. One month (26-33 days) after the injection, the rats were euthanised by decapitation, and the brains rapidly removed and frozen on dry ice.

**Figure 1:**
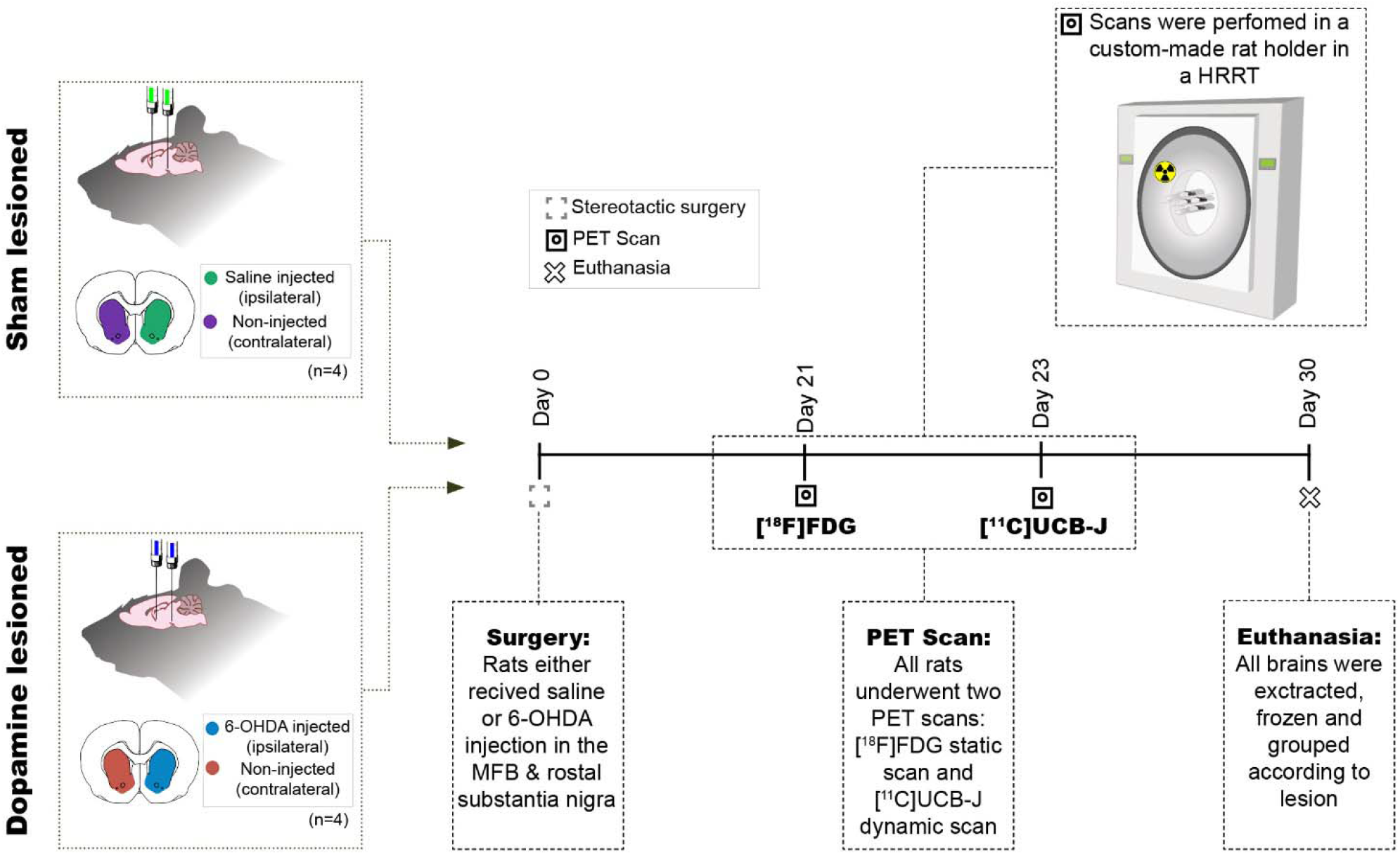
Study design. Eight rats received two intracranial injections of either 6-OHDA or saline in the medial forebrain bundle (MFB) and rostral to substantia nigra and hence divided into two groups sham lesioned (saline) or dopamine lesioned (6-OHDA). Approximately 21 days after the injections, all rats underwent an [^18^F]FDG PET scan followed by [^11^C]UCB-J PET scan 2 days after in a Siemen’s high-resolution research tomography (HRRT). All animals were euthanised 30 days after the intracranial injection.

To validate the extent of the lesion, tyrosine hydroxylase (TH) immunostaining was performed on 20 µm coronal cryosections containing the striatum. Frozen brains were sectioned on a cryostat (Leica CM1800, Leica Biosystems, Buffalo Grove, IL, USA) and mounted on Superfrost Plus(™) adhesion microscope slides (Thermo Fischer Scientific, MS, USA). Sections were stored at -80°C for the remaining period of the study. The sections were dried and processed for standard TH immunohistochemistry. Briefly, the frozen sections were first fixed in cold (4°C) 4% formaldehyde for 15 minutes. The sections were then prewashed in 0.05 M phosphate-buffered saline (PBS, pH 7.4) with 1% bovine serum albumin and then incubated overnight in a purified antiserum against TH generated in rabbits (Sigma-Aldrich, Søborg, Denmark; cat#AB152) diluted 1:500 in PBS + 0.1% Triton-X overnight at 4°C. The immunoreactivity was detected using the avidin-biotin detection method (biotinylated donkey-anti rabbit IgG (Sigma-Aldrich, Søborg, Denmark, #SAB3700966); avidin-biotin-peroxidase complex (Thermo Fischer Scientific, MS, USA #32020)) and reacted for peroxidase activity in 0.1% diamino-benzidine mixed with 0.003% H_2_O_2_ in PBS for 15 minutes. Finally, the sections were washed in distilled water and embedded in Pertex.

The stained slides were imaged on a Zeiss Axio Observer 7 using an EC Plan-Neofluoar 5x/0.16 objective by stitching multiple fields of view to cover the entire section. The resulting colour image was analysed in ImageJ 1.53G (NIH Image, Bethesda, MD, USA) by a workflow involving masking potential artefacts by automatic threshold (Moment) and conversion to 16-bit grayscale. From these, crude regions of interest encompassing the striatum were identified for quantification. Automated thresholds were used to measure the intensities in mean grey values (Minimum) and stained areas in pixel values (Moment). The intensities and areas of the ipsilateral striatum were normalised to the contralateral striatum and are presented as percentages.

### 2.4 [^18^F]FDG and [^11^C]UCB-J PET scans

All scans were performed on the Siemens HRRT (High-Resolution Research Tomography), and all rats were examined using both [^11^C]UCB-J and [^18^F]FDG. The rats were transported to the scanner at least 2 hours before the scan. Anaesthesia was induced using 3% isoflurane in oxygen. All rats were placed in a 2 × 2 custom made rat holder (illustration in Figure 1), enabling simultaneous scanning of four rats (Keller et al., 2017; Shalgunov et al., 2020; Casado-Sainz et al., 2021). While in the custom-made rat holder, the rats were kept under anaesthesia with a constant flow of isoflurane (∼2% isoflurane in oxygen). They were placed in the HRRT scanner for the time of the scan. The rats were kept warm using an infrared lamp and monitored for respiration throughout the entire scan. A rotating point source ^137^Cs transmission scan (Keller et al., 2013) was carried out before or after each emission scan.

[^18^F]FDG was acquired from the in-house clinical production of the department of clinical physiology, nuclear medicine and PET, Rigshospitalet, Denmark. Rats were fasted overnight before the scan. The animals were briefly anaesthetised, and [^18^F]FDG was administered intraperitoneal with an average injected dose of 25.05 ±3.1 MBq. The rats were placed back in their home cage to wake up from the anaesthesia to achieve [^18^F]FDG uptake while awake. Forty-five minutes after the [^18^F]FDG injection, the rats were anaesthetised, placed in the holder, and a PET emission scan was acquired for 45 minutes.

[^11^C]UCB-J was produced in-house using a modified protocol (see supplementary information) adapted from Nabulsi et al.(Nabulsi et al., 2016). The tail veins were canulated (BD Neoflon 25G, Stockholm, Sweden) before the scan. At the start of the scan, intravenous (IV) injections were given over 7-10 seconds through the tail vein catheter, with an average dose of 20.8 ± 2.1 MBq (injected mass= 0.04 ± 0.01 µg). Heparinised saline (500-600 µL) was flushed through the catheter after tracer injection. The acquisition time for [^11^C]UCB-J was 60 minutes.

### 2.5 PET image reconstruction

All list-mode data was dynamically reconstructed using ordinary Poisson 3D ordered subset expectation maximisation with point spread function modelling, resulting in PET image frames consisting of 207 planes of 256 × 256 voxels (1.22 × 1.22 × 1.22 mm). The reconstruction of the attenuation map from the transmission scan was performed using the maximum a posteriori algorithm for transmission data. All [^11^C]UCB-J scans were transformed into 33 dynamic frames (6 × 10, 6 × 20, 6 × 60, 8 × 120 and 7 × 300 seconds), while [^18^F]FDG scans were transformed into 5-minute frames and then averaged into a single frame.

### 2.6 Quantification of PET data

Pre-processing of all PET scans were done with PMOD 3.7 (PMOD Technologies, Zürich, Switzerland). Kinetic modelling was done with PMOD 3.0 (PMOD Technologies, Zürich, Switzerland). All rats were scanned in full-body, and brains were manually cropped out. For [^18^F]FDG scans, static images were manually co-registered to a standard [^18^F]FDG PET template. For [^11^C]UCB-J scans, a summed image of the last 13 frames were manually co-registered to an average T1-weighted magnetic resonance brain image in standard space. MR template used was a summed image from various rats, not part of this study, generously provided by Kristian Nygaard Mortensen. Volumes of interest (VOIs)-atlas of selected regions from the CSTC circuit from Schiffer’s atlas (Schiffer et al., 2006) were applied to the PET image in standard space. The regions (depicted in Figure 3 and Supplementary Figure 4) included in this study were: anterior cingulate cortex, medial prefrontal cortex, motor cortex, nucleus accumbens, orbitofrontal cortex, striatum, thalamus, and ventral midbrain (a region covering both the ventral tegmental area and substantia nigra). The dorsomedial striatum and dorsolateral striatum were manually delineated and used in the study (Shalgunov et al., 2020; Casado-Sainz et al., 2021). All images and co-registration were visually checked for accuracy following spatial transformation.

For [^18^F]FDG, the unit of measurement (Bq/mL) for each cropped image was transformed into standardised uptake values (SUV) by adjusting for body weight and injected dose. A whole-brain normalisation factor (WB_NF_) was calculated for each rat using [Eq. 1]. The SUV values from all the VOIs were normalised using WB_NF_.

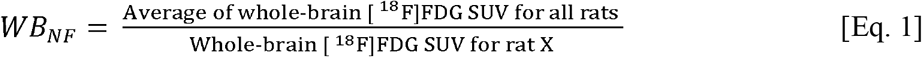

For [^11^C]UCB-J, time-activity curves (TACs) for all VOIs were extracted from the PET images. Estimates for the total blood activity was acquired using a non-invasive image-derived input function (IDIF) that was used for estimating a surrogate of V_T._ V_T_ was determined in each VOI, using the one-tissue compartment model (1TCM), which has previously been validated for [^11^C]UCB-J in mice (Bertoglio et al., 2020; Xiong et al., 2021). The IDIF was extracted from each PET image by delineating the whole blood activity in the lumen of the heart’s left ventricle. This delineation was achieved by using the ‘region growing’ function in PMOD in the early time frame by dropping a ‘seed’ at the point of highest activity in the heart and producing a VOI which is about the size of the rat’s left ventricle (5-6 voxels). In order to fit the 1TCM to the TACs, the blood volume fraction (V_B_) was fixed at 5%. In addition to V_T_, the micro-parameters K_1_ and k_2_ were also extracted from the kinetic modelling. These micro-parameters were checked for the difference due to the surgical procedure or any other reason. 1TC model fit to a representative region, ipsilateral and contralateral striatum, are shown in Supplementary Figure 5. All micro parameters (K1 and k2) for all regions are recorded in Supplementary Table 2. In addition to kinetic modelling, TACs were converted into SUVs. Ipsilateral and contralateral striatum and ventral midbrain (sham and dopamine lesioned) TACs were averaged for visual representation. This was performed using GraphPad Prism 9 (GraphPad Software, San Diego, CA, USA).

### 2.7 Statistics

Due to the limited sample size and the number of comparisons undertaken, the study is exploratory in nature, meaning that caution should be taken around drawing strong confirmatory conclusions from the data. As such, all p-values reported should be considered as a continuous assessment of indirect evidence against the null hypothesis of no difference between groups or hemispheres, and binary conclusions of “significant” or “not significant” within the Neyman-Pearson Null-hypothesis-significance-testing framework should be avoided.

The data were analysed using Jamovi (Version 1.6, The jamovi project (2021) [Computer Software]. Retrieved from https://www.jamovi.org) and RStudio (v. 4.0.3; *“Bunny-Wunnies Freak Out”*, R core team, Vienna, Austria). Graph-Pad Prism (v. 9.0.1; GraphPad Software, San Diego, CA, USA) was used for data visualisation. All data are presented as mean values ± standard deviation unless otherwise specified. The TH immunostaining comparison of the dopamine and sham lesion (ipsilateral side corrected to the contralateral side) was performed with an independent samples t-test (Mann-Whitney test).

To allow direct comparison of [^18^F]FDG normalised SUVs and [^11^C]UCB-J V_T_, Cohen’s dz values (a standardised measure of within-subject differences) between the ipsilateral regions and contralateral regions were calculated (Lakens, 2013). Cohen’s dz (standardised measure of between-group differences) values were used to compare the effect size measured by the two tracers. This shows the efficacy of detecting differences with the two radioligands.

To further explore and compare the different regions, the difference between the ipsilateral and contralateral side for each tracer ([^18^F]FDG and [^11^C]UCB-J) in the dopamine and sham lesioned groups was calculated in Jamovi using paired t-test without correction for multiple comparisons.

We performed tests on [^18^F]FDG normalised SUVs and [^11^C]UCB-J V_T_ between the two lesioned groups in regions outside the basal ganglia: thalamus, medial prefrontal cortex, anterior cingulate cortex, orbitofrontal cortex and motor cortex. These tests were performed using an independent samples t-test (Mann-Whitney test).

## 3 Results

### 3.1 Confirmation of lesion

Striatal TH immunostaining confirmed unilateral dopaminergic lesions in the striatum (Figure 2). We observed a 73.9% decrease (p = 0.03) in the stained area from the sham lesioned animals (97.50% ± 6.77) to the dopamine lesioned animals (23.54% ± 9.41). These observations were accompanied by a 24.68% reduction in staining intensity (p= 0.03) between sham lesioned (93.39% ± 4.73) and dopamine lesioned animals (68.72% ± 6.62).

**Figure 2:**
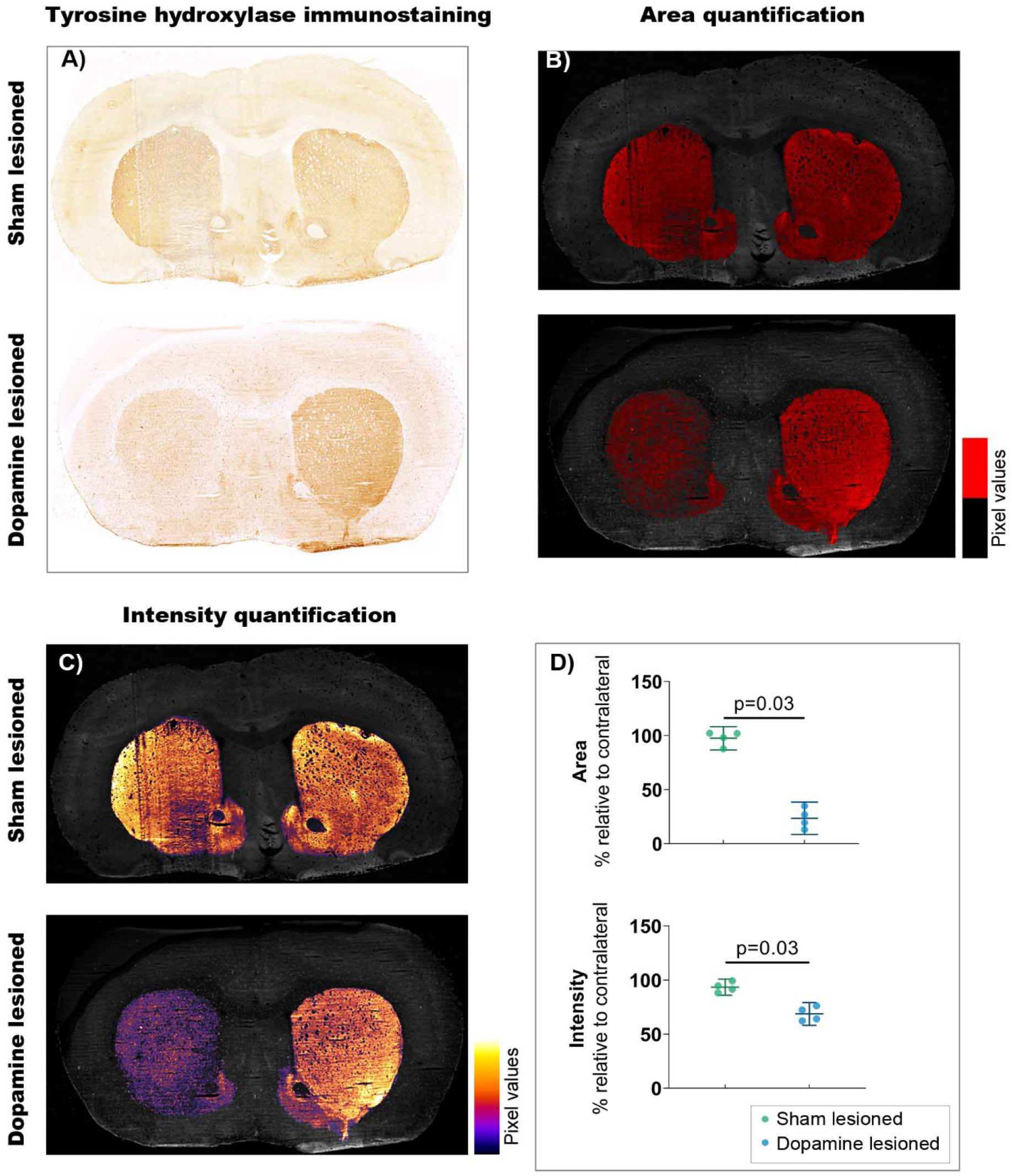
Confirmation of 6-OHDA-induced dopaminergic lesions. A) Representative example of tyrosine hydroxylate immunostaining: upper section from sham lesioned rats, lower from dopamine lesioned rats. B) Quantification of the stained area, threshold emphasised in red. C) Quantification of staining intensity, intensity scale insert. D) Quantification of intensity and area relative to contralateral striatum (n = 4/group). Error bar denotes the mean and the 95% confidence interval. P values demonstrated from the Mann-Whitney tests.

### 3.2 Representative [^11^C]UCB-J and [^18^F]FDG PET images

Representative [^11^C]UCB-J and [^18^F]FDG PET images from a rat in the dopamine and sham lesioned group are shown in Figure 3. A template structural T1 MR image is used for illustrative purpose only. Regional VOIs are shown on summed PET images in Supplementary Figure 4. For [^11^C]UCB-J, a difference was visually noticed between the ipsilateral and contralateral side of the 6-OHDA injection, especially in the striatal regions and ventral midbrain (red arrows in Figure 3). Hemispheric differences were not evident in the sham lesioned animal. For [^18^F]FDG, changes were also evident between the ipsilateral and contralateral hemisphere in the cortex, striatal regions, and ventral midbrain in the dopamine lesioned animal (red arrows in Figure 3), while no apparent differences were seen in the sham lesioned animal.

**Figure 3:**
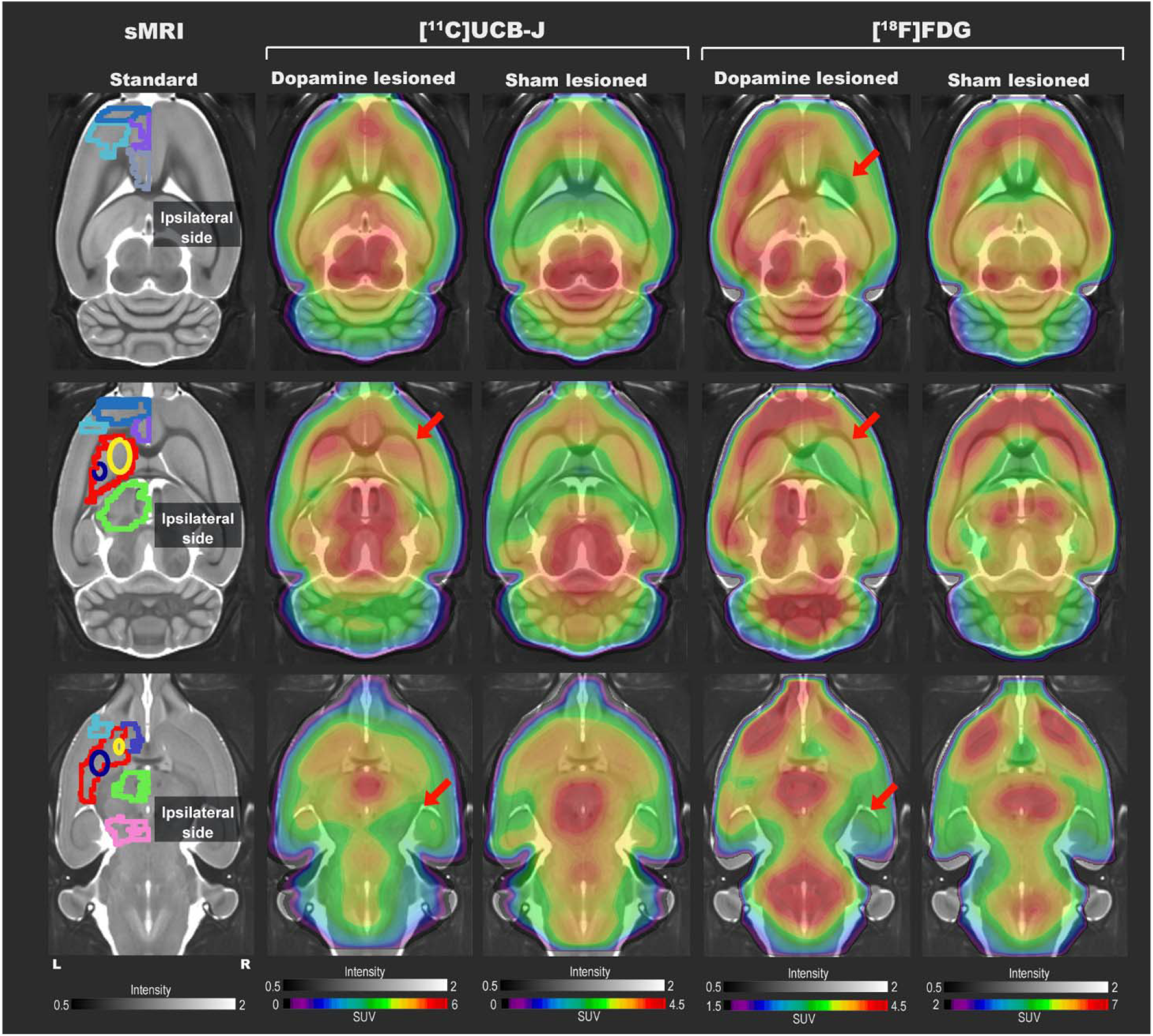
Representative [^11^C]UCB-J and [^18^F]FDG PET SUV horizontal brain slices from a dopamine and a sham lesioned rat. Standard structural MRI (for illustrative purposes) slices show the selected volumes of interest in one hemisphere; mPFC (medium blue), OFC (purple), motor cortex (light blue), ACC (grey), striatum (red), dorsomedial striatum (yellow), dorsolateral striatum (navy blue), thalamus (green), NAc(dark blue), and ventral midbrain (pink). For [^11^C]UCB-J, the SUV image represents the sum of 15-60 minutes; for [^18^F]FDG, it is the sum of all 45 minutes. The red arrow shows decreased tracer uptake in dopamine lesioned animals.

### 3.3 Decreased [^11^C]UCB-J V_T_ in dopamine lesioned hemisphere

Visually, a lower average [^11^C]UCB-J uptake can be seen through averaged TACs in the ipsilateral striatum and ventral midbrain compared to the contralateral hemisphere in dopamine lesioned animals (Figure 4 A and B). No changes were noticed in the sham lesioned animals (Figure 4 C and D). [^11^C]UCB-J V_T_ values were lower in the ipsilateral side of the striatum, dorsolateral striatum and ventral midbrain but higher in the medial prefrontal cortex and anterior cingulate cortex compared to the contralateral side (Figure 4 E and Table 1). In the sham lesioned animals, higher [^11^C]UCB-J V_T_ values were also seen in the ipsilateral anterior cingulate cortex compared to the contralateral side. No other differences were observed in [^11^C]UCB-J V_T_ (Figure 4 F and Table 1) between the ipsilateral and contralateral sides in the sham lesioned rats.

**Figure 4:**
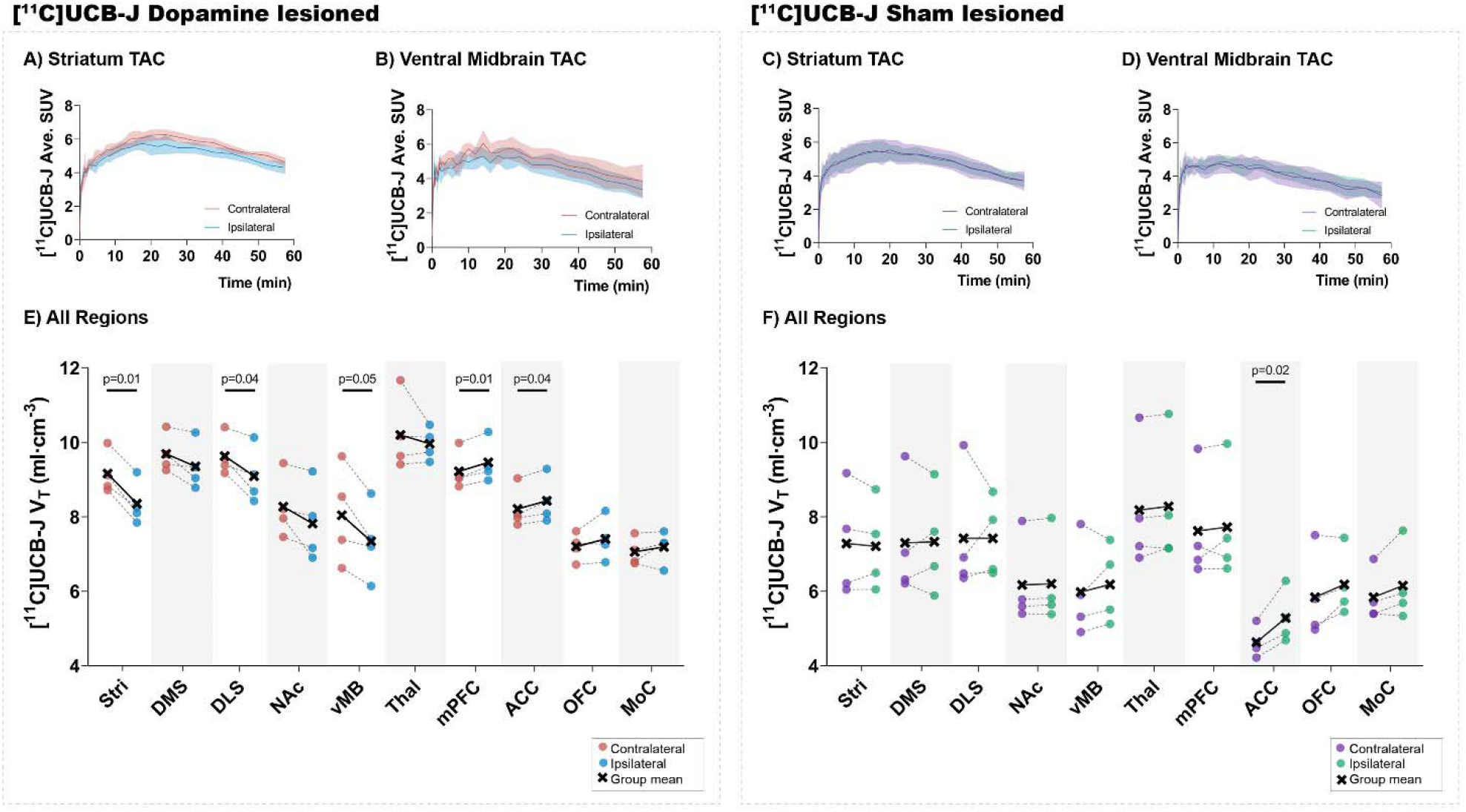
Average [^11^C]UCB-J binding. Average time-activity curves from all the animals in the striatum (A and C) and ventral midbrain (B and D). Comparison of ipsilateral and contralateral [^11^C]UCB-J V_T_ values in the selected regions of interest within the dopamine lesioned (E) and sham lesioned (F) rats. Notable differences are marked with their p values. Stri= striatum, DMS= dorsomedial striatum, DLS= dorsolateral striatum, NAc= nucleus accumbens, vMB= ventral midbrain, Thal= thalamus, mPFC= medial prefrontal cortex, ACC= anterior cingulate cortex, OFC= orbitofrontal cortex, MoC= motor cortex.

**Table 1:** Group-wise summary of the paired t-test between the ipsilateral and contralateral regions for each tracer and group. To aid overview, notable differences are marked as *. Stri= striatum, DMS= dorsomedial striatum, DLS= dorsolateral striatum, NAc= nucleus accumbens, vMB= ventral midbrain, Thal= thalamus, mPFC= medial prefrontal cortex, ACC= anterior cingulate cortex, OFC= orbitofrontal cortex, MoC= motor cortex.

| Region | [<sup>11</sup>C]UCB-J V<sub>T</sub> |  |  |  | [<sup>18</sup>F]FDG Normalized SUVs |  |  |  |
| --- | --- | --- | --- | --- | --- | --- | --- | --- |
|  | Dopamine Lesioned |  | Sham Lesioned |  | Dopamine Lesioned |  | Sham Lesioned |  |
|  | % diff | p value | % diff | p value | % diff | p value | % diff | p value |
| Stri | -8.86 % | 0.003* | -0.99 % | 0.66 | -5.66 % | 0.093 | 0.25 % | 0.926 |
| DMS | -3.48 % | 0.085 | 0.39 % | 0.919 | -7.42 % | 0.077 | -1.84 % | 0.553 |
| DLS | -5.58 % | 0.046* | 0.02 % | 0.998 | -6.30 % | 0.022* | -0.45 % | 0.893 |
| NAc | -5.35 % | 0.122 | 0.56 % | 0.173 | -7.26 % | 0.071 | 0.93 % | 0.62 |
| vMB | -8.72 % | 0.052 | 3.35 % | 0.486 | -2.89 % | 0.343 | 0.18 % | 0.821 |
| Thal | -2.55 % | 0.465 | 1.20 % | 0.233 | -4.11 % | 0.013* | 1.09 % | 0.425 |
| mPFC | 2.59 % | 0.009* | 1.33 % | 0.621 | -2.02 % | 0.147 | 1.25 % | 0.47 |
| ACC | 2.62 % | 0.043* | 14.08 % | 0.023* | -0.23 % | 0.832 | 0.47 % | 0.757 |
| OFC | 2.88 % | 0.209 | 5.71 % | 0.141 | -6.32 % | 0.020* | 0.45 % | 0.67 |
| MoC | 1.85 % | 0.435 | 5.27 % | 0.169 | -1.32 % | 0.407 | 3.25 % | 0.223 |

### 3.4 Decreased [^18^F]FDG uptake in dopamine lesioned hemisphere

There was a lower uptake of [^18^F]FDG in all striatal regions (only statistically significant in dorsolateral striatum), thalamus and orbitofrontal cortex in the ipsilateral side of dopamine lesioned rats, compared to the contralateral side (Figure 5 and Table 1). No substantial differences were found between the ipsilateral and contralateral sides within the sham lesioned animals.

**Figure 5:**
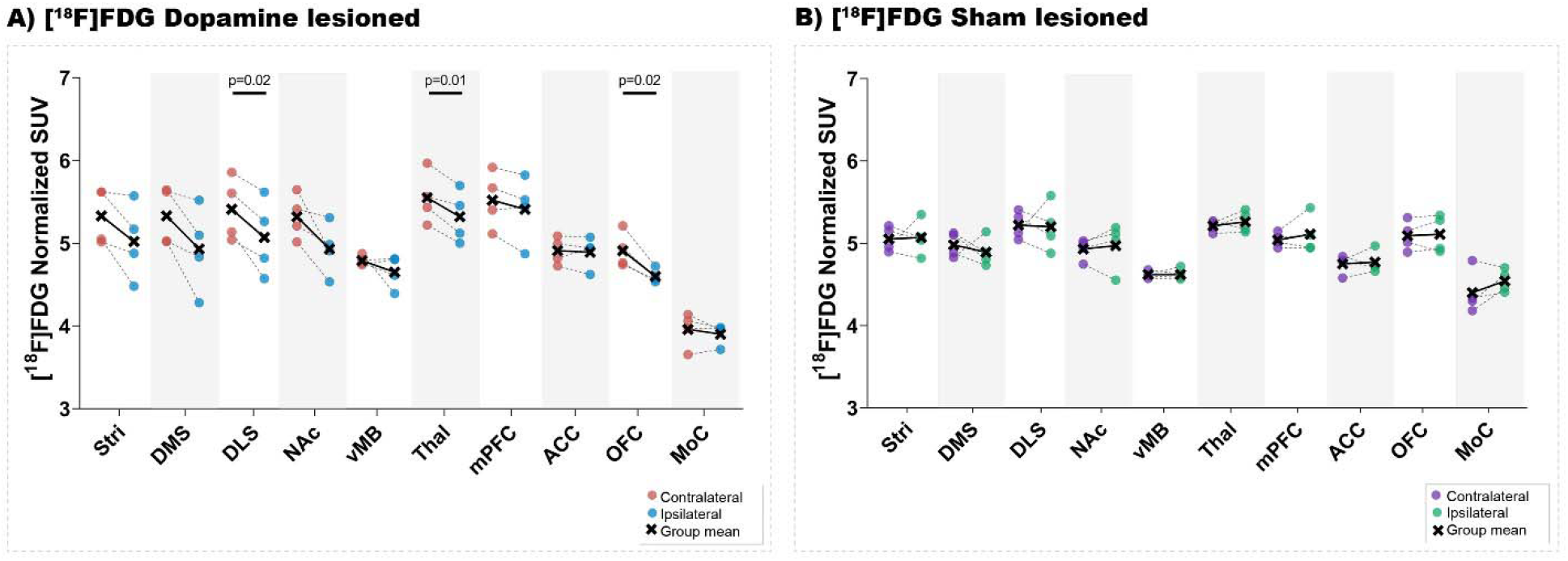
[^18^F]FDG uptake. Direct comparison between ipsilateral and contralateral hemispheres of normalised [^18^F]FDG uptake in all regions of interest within the dopamine lesioned (A) and sham lesioned (B) rats. Notable differences were marked with their uncorrected p values. Stri = striatum, DMS = dorsomedial striatum, DLS = dorsolateral striatum, NAc = nucleus accumbens, vMB = ventral midbrain, Thal = thalamus, mPFC = medial prefrontal cortex, ACC = anterior cingulate cortex, OFC = orbitofrontal cortex, MoC = motor cortex.

### 3.5 [^11^C]UCB-J and [^18^F]FDG show divergent effect sizes in dopamine and sham lesioned animals

Both [^11^C]UCB-J and [^18^F]FDG show an expected negative effect of the dopaminergic lesion in all dopamine rich regions, including the ventral midbrain, striatum, dorsomedial striatum, dorsolateral striatum and nucleus accumbens (Figure 6). Results are reported as Cohen’s dz values, showing the within-subject effect size between the ipsilateral and contralateral hemispheres. The ventral midbrain and striatum show a larger effect with [^11^C]UCB-J than [^18^F]FDG, although with confidence intervals overlapping the mean of the other radioligand. The dorsomedial striatum, dorsolateral striatum and nucleus accumbens also have overlapping confidence intervals and shows a similar effect with [^11^C]UCB-J or [^18^F]FDG.

**Figure 6:**
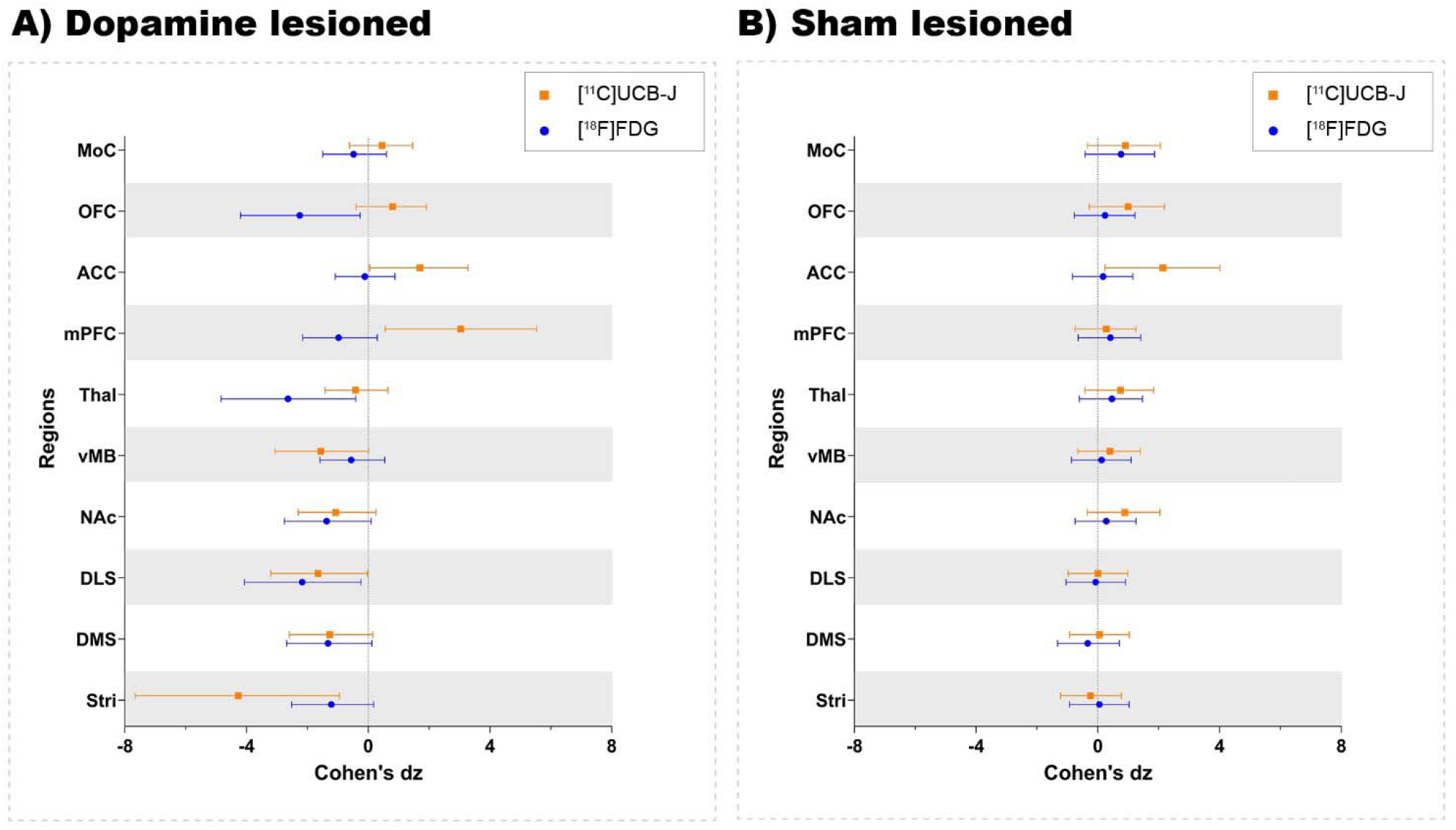
Direct comparison of effect size (Cohen’s dz values) as measured by [^11^C]UCB-J and [^18^F]FDG PET. All regions within the dopamine and sham lesioned animals in the study are compared. Error bar denotes the mean and the 95% confidence interval. Stri= striatum, DMS= dorsomedial striatum, DLS= dorsolateral striatum, NAc= nucleus accumbens, vMB= ventral midbrain, Thal= thalamus, mPFC= medial prefrontal cortex, ACC= anterior cingulate cortex, OFC= orbitofrontal cortex, MoC= motor cortex.

Besides dopamine rich regions, there is a seemingly larger reduction with [^18^F]FDG compared to [^11^C]UCB-J in the thalamus; however, the [^18^F]FDG confidence interval still includes the mean of [^11^C]UCB-J. Divergent changes can be seen in cortical regions when comparing [^11^C]UCB-J and [^18^F]FDG except for the motor cortex, which shows no effect of the dopamine lesion. In particular, the medial prefrontal cortex and orbitofrontal cortex shows a negative effect with [^18^F]FDG (higher SUV on the lesioned side), while it shows a positive effect with [^11^C]UCB-J (lower V_T_ on the lesioned side). The anterior cingulate cortex shows no effect with [^18^F]FDG but a positive effect with [^11^C]UCB-J. Sham lesioned animals do not show differences between hemispheres, except for [^11^C]UCB-J in the anterior cingulate cortex.

### 3.6 Changes in cortical regions between [^11^C]UCB-J binding and [^18^F]FDG uptake

A post hoc analysis of changes in the cortical regions and thalamus between the lesion and sham group (Figure 7) showed an increase in [^11^C]UCB-J V_T_ values in the anterior cingulate cortex (37.36%, p = 0.03) whereas there is no difference in [^18^F]FDG uptake (2.6%, p = 0.68). On the contrary, a lower [^18^F]FDG uptake is observed in the motor cortex (−16.42%, p = 0.03) and the orbitofrontal cortex (−11.08%, p = 0.03), which is not the case for [^11^C]UCB-J V_T_ (16.8%, p = 0.34 and 19.8%, p = 0.20).

**Figure 7:**
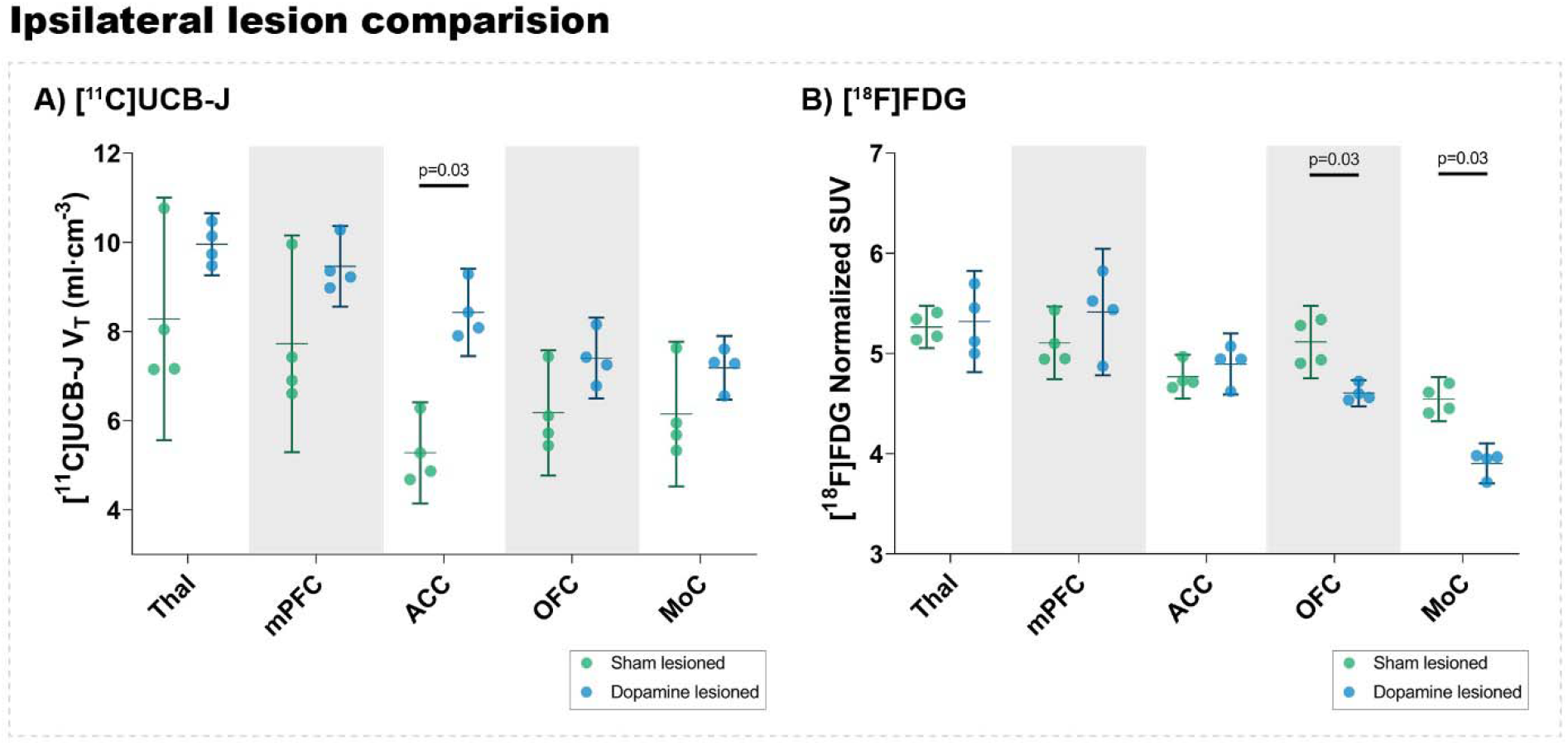
Analysis of [^11^C]UCB-J V_T_ values (A) and [^18^F]FDG (B) uptake in the ipsilateral side of dopamine lesioned and sham lesioned animals. Error bar denotes the mean and the 95% confidence interval. Thal = thalamus, mPFC = medial prefrontal cortex, ACC = anterior cingulate cortex, OFC = orbitofrontal cortex, MoC = motor cortex.

## 4 Discussion

This study explored regional differences in [^11^C]UCB-J binding and [^18^F]FDG uptake using a unilateral 6-OHDA dopaminergic lesion in rats, a commonly used animal model for PD. We observed differences in SV2A density and neuronal metabolic function between ipsilateral and contralateral hemispheres, especially the basal ganglia, which are well known to be innervated by dopaminergic terminals. This suggests a decline in dopaminergic neurons and synapses due to the 6-OHDA lesion, consistent with TH immunostaining (Figure 2).

We derived effect sizes between the ipsilateral and contralateral regions to directly compare [^11^C]UCB-J and [^18^F]FDG. The regions within the basal ganglia show similar effects with the two radioligands, lower SV2A density and metabolic function, in the ipsilateral region compared to the contralateral region. Especially lower SV2A density in the striatum, dorsolateral striatum, and ventral midbrain compared to the contralateral regions. We see a strong correlation between in vitro autoradiography ([^3^H]UCB-J fmol/mg tissue equivalent) and PET quantification ([^11^C]UCB-J V_T_) in the sham lesioned animal (Supplementary data 1.2). This further confirms the validity of the [^11^C]UCB-J PET data. A lower ipsilateral metabolic function is also observed in the regions of basal ganglia, which is consistent with previous 6-OHDA lesion studies showing an ipsilateral decrease in [^18^F]FDG uptake in the striatal regions compared to the contralateral regions (Casteels et al., 2008; Jang et al., 2012; Kordys et al., 2017). No such changes are evident in baseline animals (Supplementary data 1.1). Our observations are in line with the common understanding of the CSTC circuitry, in which the striatal response is in part sculptured by the dopaminergic input from substantia nigra (Vriend et al., 2014). Hence, diminished activity in dopamine neurons projecting to the striatum due to the 6-OHDA lesion would lead to a decline in striatal activity, as is evident from the changes in [^18^F]FDG uptake.

A difference of moderate magnitude between the ipsilateral and contralateral thalamus was noted for [^18^F]FDG but not for [^11^C]UCB-J. Although dopamine denervation of the rodent thalamus is scant (Papadopoulos and Parnavelas, 1990), we still observe decreased metabolic function. This may be due to the overall decreased function of the lesioned thalamus.

The cortical regions also show divergent group differences with [^11^C]UCB-J and [^18^F]FDG. In the orbitofrontal cortex and medial prefrontal cortex, [^18^F]FDG uptake is lower in the ipsilateral regions compared to contralateral regions. By contrast, [^11^C]UCB-J shows higher SV2A density in the ipsilateral regions compared to the contralateral regions. To our knowledge, it is the first time that a lower orbitofrontal cortex metabolic function is demonstrated in this rat model; a decrease has previously only been reported in the prefrontal cortex (Casteels et al., 2008), while other studies show unaltered metabolism (Kurachi et al., 1995). The decrease in orbitofrontal and medial prefrontal cortical metabolic function may be due to the disrupted dopaminergic innervation from the substantia nigra to the orbitofrontal cortex (Murphy and Deutch, 2018).

[^11^C]UCB-J binding is higher in the anterior cingulate cortex in most of the tests that we perform, except baseline animals (Supplementary data 1.1). While showing no effect in metabolic function, the anterior cingulate cortex’s SV2A density was higher ipsilaterally, both in the sham and dopamine lesioned animals. Likewise, the anterior cingulate cortex had higher SV2A density in the dopamine lesioned animals than sham lesioned animals, both in ipsilateral (Figure 7) and contralateral hemispheres (Supplementary Figure 3). These changes are also evident in vitro using [^3^H]UCB-J autoradiography (Supplementary data 1.2) in the sham lesioned animals (Supplementary Figure 2). Such changes in the cingulate cortex have not been previously shown in this model. We speculate that the cause is the surgery itself as the anterior cingulate cortex is part of the pain matrix (Bliss et al., 2016), but further testing is necessary to understand this observation. In addition, a reduced mechanical nociceptive threshold has been extensively reported in the 6-OHDA model, which maybe is directly related to changes in synaptic density in the anterior cingulate cortex (Buhidma et al., 2020).

We observed a lower metabolic function in the ipsilateral motor cortex and the orbitofrontal cortex between the 6-OHDA-injected and saline-injected cortexes. The difference in the motor cortex is also seen in patients with PD, but reduced metabolic function in the orbitofrontal cortex are not commonly seen in PD subjects (Meyer et al., 2017). Such cortical reduction was not detected with [^11^C]UCB-J, implying the relative robustness in detecting circuit changes with [^18^F]FDG.

Disease-specific changes in SV2A density, i.e. synaptic loss, have now been demonstrated in rodent models of neurodegeneration with intracranial injections of neurotoxic agents or with protein inoculation models of PD (Thomsen et al., 2021b, 2021a). Such synaptic loss is also demonstrated in other Alzheimer’s disease and PD mice models (Toyonaga et al., 2019; Xiong et al., 2021). Our study supports the recent study’s findings with lower SV2A density within the basal ganglia in the 6-OHDA rat model (Thomsen et al., 2021b), although there are methodological differences, such as employing different kinetic models and site of injection.

[^11^C]UCB-J has now been used in monkeys(Nabulsi et al., 2016), pigs(Thomsen et al., 2020), mice(Bertoglio et al., 2020), rats(Thomsen et al., 2021b) and humans(Finnema et al., 2016) and show favourable brain penetration, fast uptake and acceptable washout kinetics. In rats and mice, various kinetic modelling was performed using an arterial blood sampling scheme or image-derived input function (IDIF) from the heart (Bertoglio et al., 2020; Glorie et al., 2020; Thomsen et al., 2021b). The 1TCM and 2TCM both work favourably with [^11^C]UCB-J using the heart as an IDIF (Bertoglio et al., 2020; Glorie et al., 2020). The use of IDIF and whole-brain normalisation allows longitudinal studies in rodents since blood sampling often is laborious and error-prone. Although most of these studies are using mice, we assume it translates well to rats.

The small sample size is a limitation of our study, making it particularly hard to conclude that there are no differences (type 2 error). For that reason, we took an exploratory approach without pre-registered predictions and without corrections for multiple testing. As such, the results should be seen as preliminary, and we caution against confirmatory conclusions from the results and encourage future replications using larger samples and a more limited selection of analyses. Further, the contralateral hemisphere may not be an ideal control region because of the inter-hemisphere anatomical connection of the basal ganglia through the pedunculopontine nucleus (Breit et al., 2008). [^18^F]FDG results must be evaluated with caution. Other factors, such as neuroinflammation due to the injection or lesion, could evoke increased regional glucose consumption, thus concealing a decreased neuronal function (Blandini et al., 2008). Crabbé et al. have shown an increase in P2×7 receptor (key mediator in neuroinflammation), as well as translocator protein (TSPO) in 6-OHDA, lesioned animals compared to sham lesioned animals using autoradiography (Crabbé et al., 2019). These changes were significant at 21 days; hence uptake of [^18^F]FDG in the ventral midbrain may be due to neuroinflammation, which is hard to differentiate using [^18^F]FDG. Our setup in a clinical high-resolution PET scanner allows for simultaneous scanning of up to four rats, which further allowed us to perform four [^11^C]scans with a single radiosynthesis. Although this saves resources and enables a more direct comparison between rats, the resolution of the HRRT is lower than other available single-subject small animal micro-PET systems. Hence, our ability to identify potentially apparent biological differences in small regions is limited due to, e.g., partial volume effects.

Regardless, we found a pattern in the regional cortical synaptic density and neuronal metabolic function, which could be clinically relevant, especially changes within the anterior cingulate cortex and orbitofrontal cortex. We see a clear advantage of including both tracers to get a clearer picture of the neuropathology of neurodegenerative diseases like PD.

## 5 Conclusion

[^11^C]UCB-J and [^18^F]FDG PET revealed similar changes in the basal ganglia following 6-OHDA dopaminergic lesion in rats. A region-based analysis suggested a divergent response to lesions, especially in the cortical regions, orbitofrontal cortex and medial prefrontal cortex, where higher synaptic density yet lower neuronal metabolic function was observed. Taken together, the results suggest that combined [^11^C]UCB-J and [^18^F]FDG scans may yield a better understanding of aberrant CSTC circuit function and a better diagnostic outcome in patients with neurodegenerative disorders.

## 7 Author Contributions

Conceptualisation, NRR, FG, PPS, MP; methodology, NRR, FG, PPS, MP; software, NRR, FG, PPS, MP; validation, NRR, FG, PPS; formal analysis, NRR, FG, MJ; investigation, NRR, FG, IVA, NRS, AV; resources, NRR, MJ, MP; data curation, NRR, MP; writing—original draft preparation, NRR.; writing—review and editing, NRR, FG, MJ, INP, PPS, GMK, MP; visualisation, NRR; supervision, JM, PMF, MMH, PPS, GMK, MP; funding acquisition, NRR, GMK, MP. All authors have read and agreed to the published version of the manuscript.

## 8 Funding

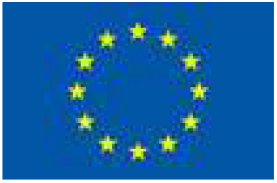

This project has received funding from the European Union’s Horizon 2020 research and innovation programme under the Marie Skłodowska-Curie grant agreement No 813528. This project also received funding from Parkinson foreningen, Denmark (R16-A247). MP received funding from the Lundbeck Foundation (R192-2015-1591 and R194-2015-1589), Augustinus Foundation (18-3746 and 17-1982), Independent Research Fund Denmark (5053-00036B), Savværksejer Jeppe Juhls og Hustrus Ovita Juhls Mindelegat and Købmand i Odense Johann og Hanne Weimann født Seedorffs Legat.

## 9 Acknowledgements

The authors would like to thank and show sincere gratitude to the veterinarians and staff at the Department of Experimental Medicine, the University of Copenhagen, for their continued assistance with animal experiments. The authors would also like to thank the Department of Clinical Physiology, Nuclear Medicine and PET for their continuous support in imaging. Structural reference rat MR image is generously provided by Kristian Nygaard Mortensen, Center for Translational Neuromedicine, University of Copenhagen.

## 11 Data Availability Statement

All data is made available at a GitHub repository (https://github.com/nakulrrraval/6-OHDA-rat-PET-paper). All other requests are directed to the corresponding or first author of this article.

## References

Bertoglio, D., Verhaeghe, J., Miranda, A., Kertesz, I., Cybulska, K., Korat, Š., et al. (2020). Validation and non-invasive kinetic modeling of [11C]UCB-J PET imaging in mice. J. Cereb. Blood Flow Metab. 40, 1351–1362. doi:10.1177/0271678X19864081.

Blandini, F., Armentero, M. T., and Martignoni, E. (2008). The 6-hydroxydopamine model: News from the past. Park. Relat. Disord. 14, S124–S129. doi:10.1016/j.parkreldis.2008.04.015.

Bliss, T. V. P., Collingridge, G. L., Kaang, B. K., and Zhuo, M. (2016). Synaptic plasticity in the anterior cingulate cortex in acute and chronic pain. Nat. Rev. Neurosci. 17, 485–496. doi:10.1038/nrn.2016.68.

Breit, S., Martin, A., Lessmann, L., Cerkez, D., Gasser, T., and Schulz, J. B. (2008). Bilateral changes in neuronal activity of the basal ganglia in the unilateral 6-hydroxydopamine rat model. J. Neurosci. Res. 86, 1388–1396. doi:10.1002/jnr.21588.

Buhidma, Y., Rukavina, K., Chaudhuri, K. R., and Duty, S. (2020). Potential of animal models for advancing the understanding and treatment of pain in Parkinson’s disease. npj Park. Dis. 6, 1–7. doi:10.1038/s41531-019-0104-6.

Casado-Sainz, A., Gudmundsen, F., Baerentzen, S. L., Lange, D., Ringsted, A., Martinez-Tajada, I., et al. (2021). Nigro-striatal dopamine activation lowers behavioral and neuronal phenotypes associated with obsessive-compulsive disorder. bioRxiv, 2021.02.11.430770. doi:10.1101/2021.02.11.430770.

Casteels, C., Lauwers, E., Bormans, G., Baekelandt, V., and Van Laere, K. (2008). Metabolic-dopaminergic mapping of the 6-hydroxydopamine rat model for Parkinson’s disease. Eur. J. Nucl. Med. Mol. Imaging 35, 124–134. doi:10.1007/s00259-007-0558-3.

Crabbé, M., Van Der Perren, A., Bollaerts, I., Kounelis, S., Baekelandt, V., Bormans, G., et al. (2019). Increased P2×7 receptor binding is associated with neuroinflammation in acute but not chronic rodent models for Parkinson’s disease. Front. Neurosci. 13. doi:10.3389/fnins.2019.00799.

Esteban, S., Lladó, J., Sastre-Coll, A., and García-Sevilla, J. A. (1999). Activation and desensitization by cyclic antidepressant drugs of α2-autoreceptors, α2-heteroreceptors and 5-HT(1A)-autoreceptors regulating monoamine synthesis in the rat brain in vivo. Naunyn. Schmiedebergs. Arch. Pharmacol. 360, 135–143. doi:10.1007/s002109900045.

Finnema, S. J., Nabulsi, N. B., Eid, T., Detyniecki, K., Lin, S. F., Chen, M. K., et al. (2016). Imaging synaptic density in the living human brain. Sci. Transl. Med. 8. doi:10.1126/scitranslmed.aaf6667.

Glorie, D., Verhaeghe, J., Miranda, A., De Lombaerde, S., Stroobants, S., and Staelens, S. (2020). Sapap3 deletion causes dynamic synaptic density abnormalities: a longitudinal [11C]UCB-J PET study in a model of obsessive–compulsive disorder-like behaviour. EJNMMI Res. 10. doi:10.1186/s13550-020-00721-2.

Jang, D. P., Min, H. K., Lee, S. Y., Kim, I. Y., Park, H. W., Im, Y. H., et al. (2012). Functional neuroimaging of the 6-OHDA lesion rat model of Parkinson’s disease. Neurosci. Lett. 513, 187–192. doi:10.1016/j.neulet.2012.02.034.

Keller, S. H., L’Estrade, E. N., Dall, B., Palner, M., and Herth, M. (2017). Quantification accuracy of a new HRRT high throughput rat hotel using transmission-based attenuation correction: A phantom study. in 2016 IEEE Nuclear Science Symposium, Medical Imaging Conference and Room-Temperature Semiconductor Detector Workshop, NSS/MIC/RTSD 2016 (IEEE), 1–3. doi:10.1109/NSSMIC.2016.8069467.

Keller, S. H., Svarer, C., and Sibomana, M. (2013). Attenuation correction for the HRRT PET-scanner using transmission scatter correction and total variation regularization. IEEE Trans. Med. Imaging 32, 1611–1621. doi:10.1109/TMI.2013.2261313.

Kordys, E., Apetz, N., Schneider, K., Duncan, E., Büschbell, B., Rohleder, C., et al. (2017). Motor impairment and compensation in a hemiparkinsonian rat model: correlation between dopamine depletion severity, cerebral metabolism and gait patterns. EJNMMI Res. 7. doi:10.1186/s13550-017-0317-9.

Kurachi, M., Yasui, S. I., Kurachi, T., Shibata, R., Murata, M., Hagino, H., et al. (1995). Hypofrontality does not occur with 6-hydroxydopamine lesions of the medial prefrontal cortex in rat brain. Eur. Neuropsychopharmacol. 5, 63–68. doi:10.1016/0924-977X(94)00136-Y.

Lakens, D. (2013). Calculating and reporting effect sizes to facilitate cumulative science: A practical primer for t-tests and ANOVAs. Front. Psychol. 4, 863. doi:10.3389/fpsyg.2013.00863.

Loane, C., and Politis, M. (2011). Positron emission tomography neuroimaging in Parkinson’s disease. Am. J. Transl. Res. 3, 323–341. Available at: www.ajtr.org [Accessed March 27, 2021].

Meyer, P. T., Frings, L., Rücker, G., and Hellwig, S. (2017). 18F-FDG PET in Parkinsonism: Differential diagnosis and evaluation of cognitive impairment. J. Nucl. Med. 58, 1888–1898. doi:10.2967/jnumed.116.186403.

Murphy, M. J. M., and Deutch, A. Y. (2018). Organization of afferents to the orbitofrontal cortex in the rat. J. Comp. Neurol. 526, 1498–1526. doi:10.1002/cne.24424.

Nabulsi, N. B., Mercier, J., Holden, D., Carr, S., Najafzadeh, S., Vandergeten, M. C., et al. (2016). Synthesis and preclinical evaluation of 11C-UCB-J as a PET tracer for imaging the synaptic vesicle glycoprotein 2A in the brain. J. Nucl. Med. 57, 777–784. doi:10.2967/jnumed.115.168179.

Papadopoulos, G. C., and Parnavelas, J. G. (1990). Distribution and synaptic organization of dopaminergic axons in the lateral geniculate nucleus of the rat. J. Comp. Neurol. 294, 356–361. doi:10.1002/cne.902940305.

Schiffer, W. K., Mirrione, M. M., Biegon, A., Alexoff, D. L., Patel, V., and Dewey, S. L. (2006). Serial microPET measures of the metabolic reaction to a microdialysis probe implant. J. Neurosci. Methods 155, 272–284. doi:10.1016/j.jneumeth.2006.01.027.

Shalgunov, V., Xiong, M., L’Estrade, E. T., Raval, N. R., Andersen, I. V., Edgar, F. G., et al. (2020). Blocking of efflux transporters in rats improves translational validation of brain radioligands. EJNMMI Res. 10, 124. doi:10.1186/s13550-020-00718-x.

Thomsen, M. B., Ferreira, S. A., Schacht, A. C., Jacobsen, J., Simonsen, M., Betzer, C., et al. (2021a). PET imaging reveals early and progressive dopaminergic deficits after intra-striatal injection of preformed alpha-synuclein fibrils in rats. Neurobiol. Dis. 149. doi:10.1016/j.nbd.2020.105229.

Thomsen, M. B., Jacobsen, J., Lillethorup, T. P., Schacht, A. C., Simonsen, M., Romero-Ramos, M., et al. (2021b). In vivo imaging of synaptic SV2A protein density in healthy and striatal-lesioned rats with [11C]UCB-J PET. J. Cereb. Blood Flow Metab. 41, 819–830. doi:10.1177/0271678X20931140.

Thomsen, M. B., Schacht, A. C., Alstrup, A. K. O., Jacobsen, J., Lillethorup, T. P., Bærentzen, S. L., et al. (2020). Preclinical PET Studies of [11C]UCB-J Binding in Minipig Brain. Mol. Imaging Biol. 22, 1290–1300. doi:10.1007/s11307-020-01506-8.

Toyonaga, T., Smith, L. M., Finnema, S. J., Gallezot, J. D., Naganawa, M., Bini, J., et al. (2019). In vivo synaptic density imaging with 11C-UCB-J detects treatment effects of saracatinib in a mouse model of Alzheimer disease. J. Nucl. Med. 60, 1780–1786. doi:10.2967/jnumed.118.223867.

Vriend, C., Pattij, T., Van Der Werf, Y. D., Voorn, P., Booij, J., Rutten, S., et al. (2014). Depression and impulse control disorders in Parkinson’s disease: Two sides of the same coin? Neurosci. Biobehav. Rev. 38, 60–71. doi:10.1016/j.neubiorev.2013.11.001.

Xiong, M., Roshanbin, S., Rokka, J., Schlein, E., Ingelsson, M., Sehlin, D., et al. (2021). In vivo imaging of synaptic density with [11C]UCB-J PET in two mouse models of neurodegenerative disease. Neuroimage 239, 118302. doi:10.1016/j.neuroimage.2021.118302.

