## Supplemental data and figures v3 for "Synaptic density and neuronal metabolic function measured by PET in the unilateral 6-OHDA rat model of Parkinson’s disease"

Supplementary Material

### 1 Supplementary Data

#### 1.1 Baseline [^11^C]UCB-J

##### 1.1.1 Methods and Materials:

To validate any potential hemisphere asymmetry we acquired baseline [^11^C]UCB-J PET in an additional group of eight female Long-Evans WT rats (193 ± 6 g, 9-10 weeks old when scanned) (Janvier). The animals were held under standard laboratory conditions with 12-hour light/12-hour dark cycles and ad libitum access to food and water. All animal experiments conformed to the European Commission’s Directive 2010/63/EU with approval from the Danish Council of Animal Ethics (Journal no. 2017-15-0201-01283) and the Department of Experimental Medicine, University of Copenhagen.

Scans were performed in the same fashion as previously described in the main manuscript. Briefly, after transport to the Siemens HRRT scanner, anaesthesia was induced using 3% isoflurane in oxygen. All rats were placed in a 2 x 2 custom made rat holder. Four rats were scanned at a time. While in the custom-made rat holder, the rats were kept under anaesthesia with a constant flow of isoflurane (~2% isoflurane in oxygen). At the start of the scan, intravenous (IV) injections were given over 7-10 secs through the tail vein catheter, with an average dose of 17.2 ± 2.5 MBq (injected mass= 0.06 ± 0.07 µg). Heparinised saline (500-600 µL) was flushed through the catheter after tracer injection. The acquisition time for [^11^C]UCB-J was 60 minutes.

PET image reconstruction and pre-processing were also performed in the same fashion as previously described. The data were analysed using Jamovi (Version 1.6, The jamovi project (2021) [Computer Software]. The difference between the left and right hemispheres for [^11^C]UCB-J in the baseline animals was calculated in Jamovi using paired t-test without correction for multiple comparisons. Cohen’s dz values between the regions in the right and left hemispheres were also calculated. Graph-Pad Prism (v. 9.2.0; GraphPad Software, San Diego, CA, USA) was used for data visualisation.

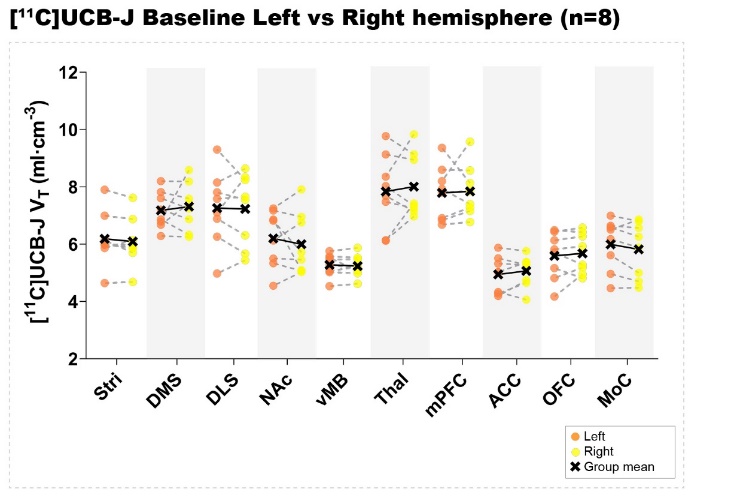

**Supplementary Figure 1.** Baseline [^11^C]UCB-J. Comparison of the [^11^C]UCB-J V_T_ between the left and right hemisphere in baseline animals (n=8). Stri= striatum, DMS= dorsomedial striatum, DLS= dorsolateral striatum, NAc= nucleus accumbens, vMB= ventral midbrain, Thal= thalamus, mPFC= medial prefrontal cortex, ACC= anterior cingulate cortex, OFC= orbitofrontal cortex, MoC= motor cortex.

**Supplementary Table 1.** Summary of the paired t-test between the regions in the left and right hemisphere for [^11^C]UCB-J in the baseline animals (n=8). Cohen’s dz values and it’s 95% cofindence interval has also been added. Stri= striatum, DMS= dorsomedial striatum, DLS= dorsolateral striatum, NAc= nucleus accumbens, vMB= ventral midbrain, Thal= thalamus, mPFC= medial prefrontal cortex, ACC= anterior cingulate cortex, OFC= orbitofrontal cortex, MoC= motor cortex.

|  | **[^11^C]UCB-J V_T_ in left vs right hemisphere** | | | | |
| --- | --- | --- | --- | --- | --- |
|  | Baseline animals | | | | |
| **Region** | **% diff** | **p value** | **Cohen’s dz** | **Cohen’s dz 95% CI** | |
|  |  |  |  | **Upper** | **Lower** |
| **Stri** | 1.50% | 0.254 | -0.4394 | -1.154 | 0.303 |
| **DMS** | -1.82% | 0.649 | 0.168 | -0.536 | 0.861 |
| **DLS** | 0.41% | 0.913 | -0.04 | -0.732 | 0.655 |
| **NAc** | 3.26% | 0.528 | -0.235 | -0.93 | 0.476 |
| **vMB** | 0.82% | 0.553 | -0.2203 | -0.915 | 0.489 |
| **Thal** | -2.16% | 0.61 | 0.1889 | -0.517 | 0.882 |
| **mPFC** | -0.69% | 0.844 | 0.072 | -0.624 | 0.763 |
| **ACC** | -2.45% | 0.317 | 0.3808 | -0.352 | 1.089 |
| **OFC** | -1.52% | 0.552 | 0.2208 | -0.489 | 0.915 |
| **MoC** | 2.91% | 0.126 | -0.615 | -1.359 | 0.164 |

##### 1.1.2 Results:

No differences were observed in [^11^C]UCB-J V_T_ (Supplementary Figure 1 and Supplementary Table 1) between the regions in the left and right hemispheres at baseline.

#### 1.2 Autoradiography with [^3^H]UCB-J

##### 1.2.1 Methods and Materials:

Autoradiography was performed using [^3^H]UCB-J (Pharmaron Ltd., Hoddesdon, UK, molar activity 28 Ci/mmol). Radio-Thin-Layered-Chromatography (R-TLC) was performed to measure the radiochemical purity (RCP) and integrity of the parent compound. The mobile phase for [^3^H]UCB-J R-TLC was Acetonitrile:Ammonium formate [25:75] (0.1 M, with 0.5% AcOH, pH 4.2). Frozen sections (containing three slices) from sham lesioned animals (n=4) (20 µm, sectioning previously explained) were thawed to room temperature for 30–45 min before preincubating twice for 10 min in 50 mM Tris-HCl pre-incubation buffer set to 7.4 pH containing 0.5% bovine serum albumin (BSA). The sections were incubated in assay buffer containing 6 nM [^3^H]UCB-J in 50 mMTris-HCl buffer containing 5 mM MgCl2, 2 mM EGTA and 0.5% BSA (pH 7.4) for 1 hour. Incubation was terminated by two 10-min washes with ice-cold pre-incubation buffer followed by a rapid rinse in ice-cold deionised H_2_O (dH_2_O). After washing, the slides were rapidly air-dried and fixated in a paraformaldehyde vapour chamber overnight in cold storage (4 °C). The next day, the samples were moved to an exicator for 45–60 min to remove any excess moisture and then placed in a cassette for autoradiography with tritium sensitive image plates (BAS-IP TR2040, Science Imaging, Scandinavia AB, Nacka, Sweden) along with radioactive tritium standards (ART0123B, American Radiolabelled Chemical, Inc., St. Loui, MO, USA). The image plates were exposed for two days. After the exposure, the image plates were read using a Fujifilm BAS 1000 scanner (Fujifilm Europe, GmbH, Duesseldorf, Germany). Calibration, quantification and data evaluation was done using ImageJ software (NIH Image, Bethesda, MD, USA). The regions of interest were hand-drawn or drawn using the wand tool and visually inspected after automated delineation. The four-parameter general curve fit (David Rodbard, NIH) of decay corrected tritium standards was used to convert mean pixel density (grayscale) to nCi/mg tissue equivalent (TE). Total binding was determined in the grey matter: striatum, ventral midbrain, cingulate cortex and motor cortex (Supplementary Figure 2B). Non-specific binding was determined in the corpus callosum or the deep cortical white matter (Supplementary Figure 2B). We have previously validated the use of white matter as non-specific binding in pigs (Raval et al., 2021); we have also validated this in rats (data not shown). Finally, the decay-corrected molar activity of the representative radioligand was used to convert nCi/mg TE to fmol/mg TE. Specific binding was calculated as the difference between total binding and non-specific binding. The data were analysed using Jamovi (Version 1.6, The jamovi project (2021) [Computer Software]. Retrieved from <https://www.jamovi.org>). The difference between the ipsilateral and contralateral side [^3^H]UCB-J in the sham lesioned group was calculated in Jamovi using paired t-test without correction for multiple comparisons. Graph-Pad Prism (v. 9.2.0; GraphPad Software, San Diego, CA, USA) was used for data visualization. Graph-Pad Prism was also used to calculate Pearson r and perform a simple linear regression. Pearson r values between the [^3^H]UCB-J fmol/mg TE and representative [^11^C]UCB-J V_T_ values from each region in the contralateral and ipsilateral hemisphere providing eight different correlation values. Values were averaged for the contralateral and ipsilateral separately to get two Pearson r value.

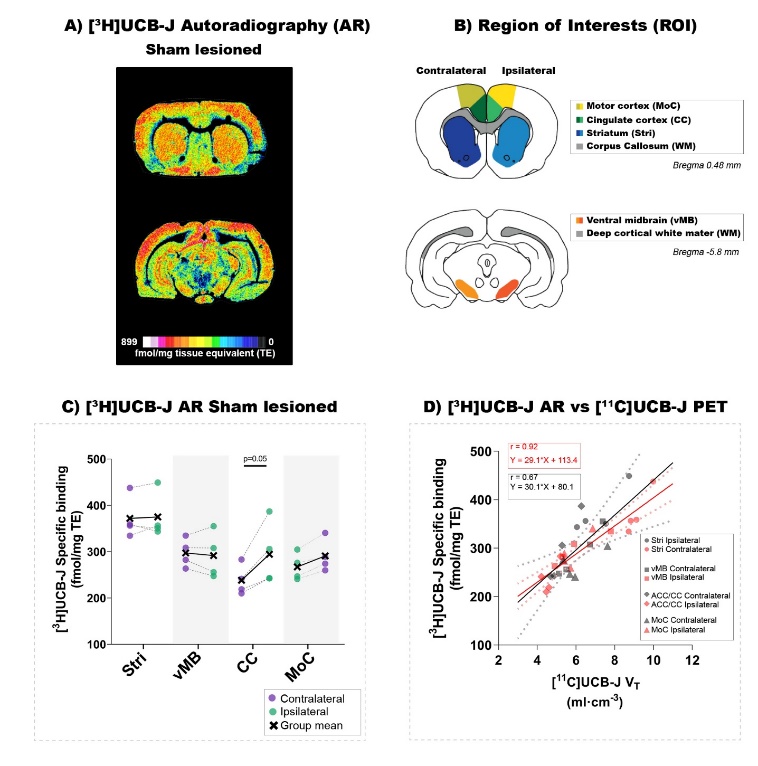

**Supplementary Figure 2.** [^3^H]UCB-J in vitro autoradiography. A) Representative examples of [^3^H]UCB-J autoradiography from sham lesioned animals: upper section contains the striatum, cingulate cortex, and motor cortex, the lower section contains the ventral midbrain. B) Region of interests used in the study includes: Stri = Striatum (blue), vMB = ventral midbrain (orange), CC = cingulate cortex (green), MoC = motor cortex (yellow). C) Comparison of ipsilateral and contralateral [^3^H]UCB-J specific binding (fmol/mg tissue equivalent [TE]) values in the selected regions of interest within the sham lesioned. Notable differences are marked with their p values. D) Correlation between the [^3^H]UCB-J fmol/mg TE and representative [^11^C]UCB-J V_T_ values in the sham lesioned animals (n=4). Average Pearson r values and equation is inserted colour coded for the two linear regression.

##### 1.2.2 Results

Representative [^3^H]UCB-J autoradiography images from sham lesioned animals are shown in Supplementary Figure 2A. Supplementary Figure 2B shows the region of interest used in this analysis. Increased specific [^3^H]UCB-J binding in the ipsilateral hemisphere compared to the contralateral hemisphere is seen in the cingulate cortex (23.3%, Cohen dz = 1.54, p = 0.05) in sham lesioned animals (Supplementary Figure 1C). A correlation analysis between [^3^H]UCB-J fmol/mg TE and representative [^11^C]UCB-J V_T_ values in both the contralateral and ipsilateral regions in the sham lesioned animals shows a Pearson r value of 0.8815.

#### 1.3 Changes in contralateral regions of [^11^C]UCB-J binding and [^18^F]FDG uptake

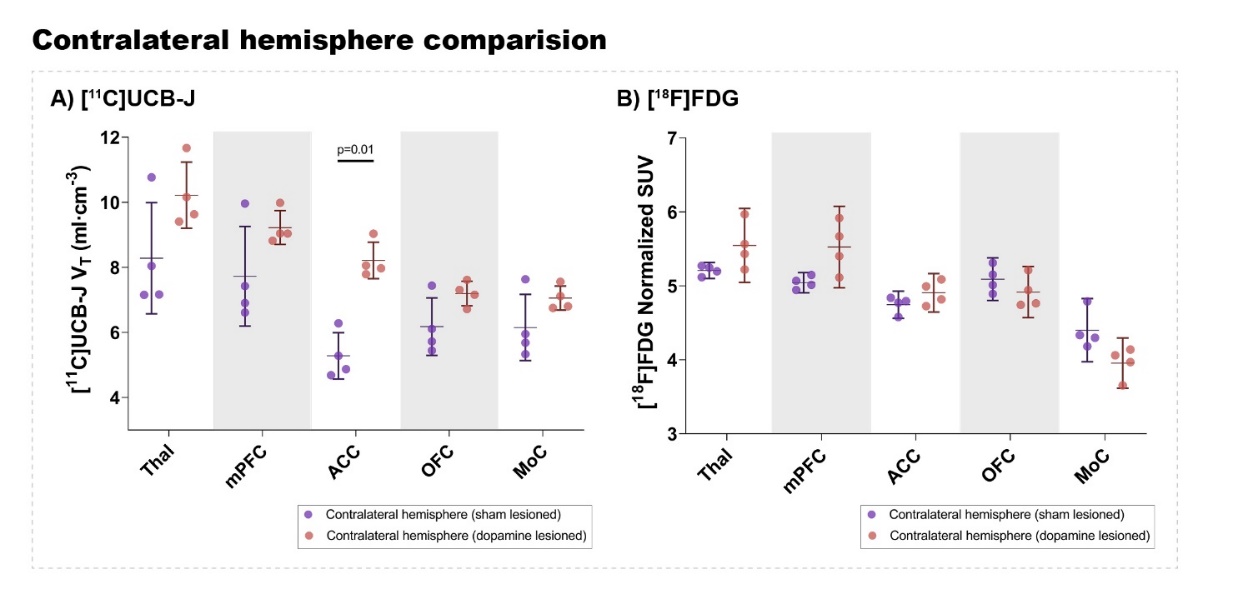

**Supplementary Figure 3.** Analysis of [^11^C]UCB-J V_T_ values (A) and [^18^F]FDG (B) uptake in the contralateral side of dopamine lesioned and sham lesioned animals. Error bar denotes the mean and the 95% confidence interval. Thal = thalamus, mPFC = medial prefrontal cortex, ACC = anterior cingulate cortex, OFC = orbitofrontal cortex, MoC = motor cortex.

A posthoc analysis of changes in the cortical regions and thalamus in the contralateral hemisphere between the dopamine lesion and sham group (Supplementary Figure 3) showed an increase in [^11^C]UCB-J V_T_ values in the anterior cingulate cortex (55.5%, p = 0.01). In contrast, there is no difference in [^18^F]FDG uptake.

#### 1.4 Radiosynthesis of [^11^C]UCB-J ((R)-1-((3-(11C-methyl-11C)pyridin-4-yl)methyl)-4-(3,4,5-trifluorophenyl)pyrrolidin-2-one)

Proton irradiation of the target material (nitrogen-14 gas) is performed using the cyclotron: Scanditronix MC-32 (Variable energy: 16-32 MeV protons) with an aluminium high-pressure gas target. Irradiations for carbon-11 are performed at 16 MeV. The target gas used is 10% hydrogen in nitrogen_._ Following irradiation, the target gas is transferred to the radiochemistry system through stainless steel capillaries.

Radiosynthesis is carried out using a fully automated radiochemistry system manufactured by Scansys Aps. [^11^C]methyl iodide is synthesised from [^11^C]methane by a standard circulation procedure.

##### 1.4.1 Preparation of the precursor:

1 M HCl (15 µL) and MeOH (70 µL) is added to the precursor (1.2 mg) and reacted overnight. On the day of synthesis, the liquid is removed by a stream of nitrogen to complete dryness.

##### 1.4.2 Preparation of the labelling mixture:

4-5 mg P-(o-tolyl)_3_ was dissolved in DMF (1.8 mL) and H_2_O (0.2 mL) in a capped vial and degassed with nitrogen. Pd_2_dba_3_ (4-5 mg) was weighed out in a capped vial and flushed with nitrogen. Right before labelling, the P-(o-tolyl)_3_ solution was added to the Pd_2_dba_3_. From here, 350 µL was withdrawn and added to a 0.9 ml vial containing 0.5 M K_2_CO_3_ (20 µL).

The formed [^11^C]methyl iodide was trapped in the 0.9 ml glass vial containing the K_2_CO_3_/ P-(o-tolyl)_3_/Pd_2_dba_3_ mixture. After trapping the [^11^C]methyl iodide, the hydrolysed DM-BF_3_-UCB-J precursor was re-dissolved in DMF (150 ul) added and reacted with the mixture by heating at 100 °C for 300 seconds to give [^11^C]UCB-J. The diluted reaction mixture was diluted with 4 ml 0.1 % H_3_PO_4_ and automatically injected onto a preparative HPLC column (Onyx™ Monolithic C-18, 100 ×10 mm equipped with a SecurityGuard Cartridge Lux Cellulose-4, 4x3.0 mm; flow: 6 ml/min; eluent: 12.5/80 [ethanol (96%)/0.1 M phosphoric acid]). The radioactive fraction corresponding to the radiolabelled product (retention time ca. 300 s) was collected by diverting the flow from the column outlet through a 0.22 µm sterile filter and directly into a sterile stoppered and a capped vial containing phosphate buffer (9 ml, pH 7). The identity and molar activity of the product was determined by using a C-18 column (Kinetex 2.6um, C18, 100Å, 50 x 4.6 mm, Phenomenex) eluted with 33% acetonitrile in 25mM citrate buffer (pH 5.4); injection volume 50 µl; flow rate 1.5 ml/min; on-line UV (261 nm) and radioactivity detection. Rt UCB-J 2.1 min.

##### 1.4.3 Molar activity:

The molar activity at the time of injection for the two productions used in this 6-OHDA study was 404 and 168 GBq/µmol. The molar activity at the time of injection in the baseline study was 353 and 628 GBq/µmol.

### 2 Supplementary Figures and Tables

#### 2.1 Supplementary Figures

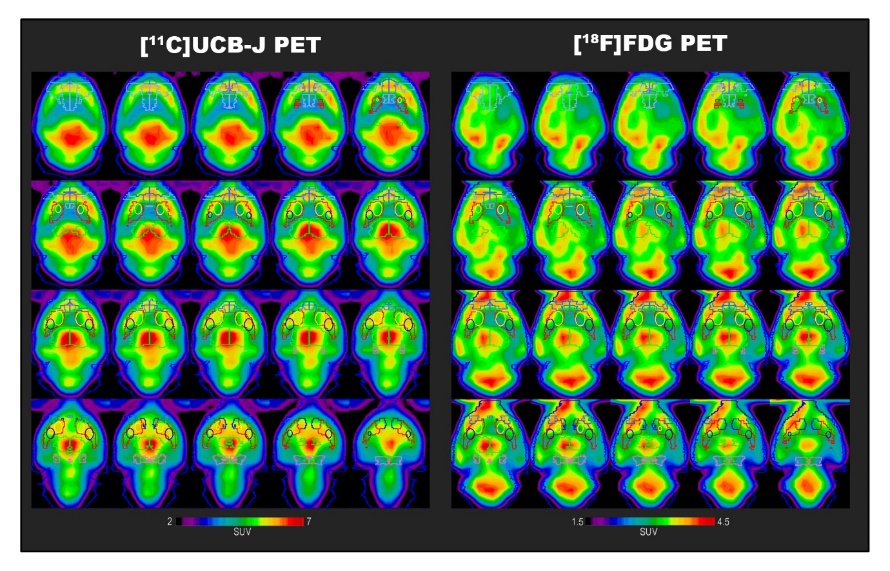

**Supplementary Figure 4.** Position and fit of select region of interests on summed PET images from the study of a single representative animal for [^18^F]FDG and [^11^C]UCB-J scan. Regions include: medial prefrontal cortex (medium blue), orbitofrontal cortex (purple), motor cortex(light blue), anterior cingulate cortex (grey), striatum (red), dorsomedial striatum (yellow), dorsolateral striatum (navy blue), thalamus (green), nucleus accumbens (dark blue), and ventral midbrain (pink).

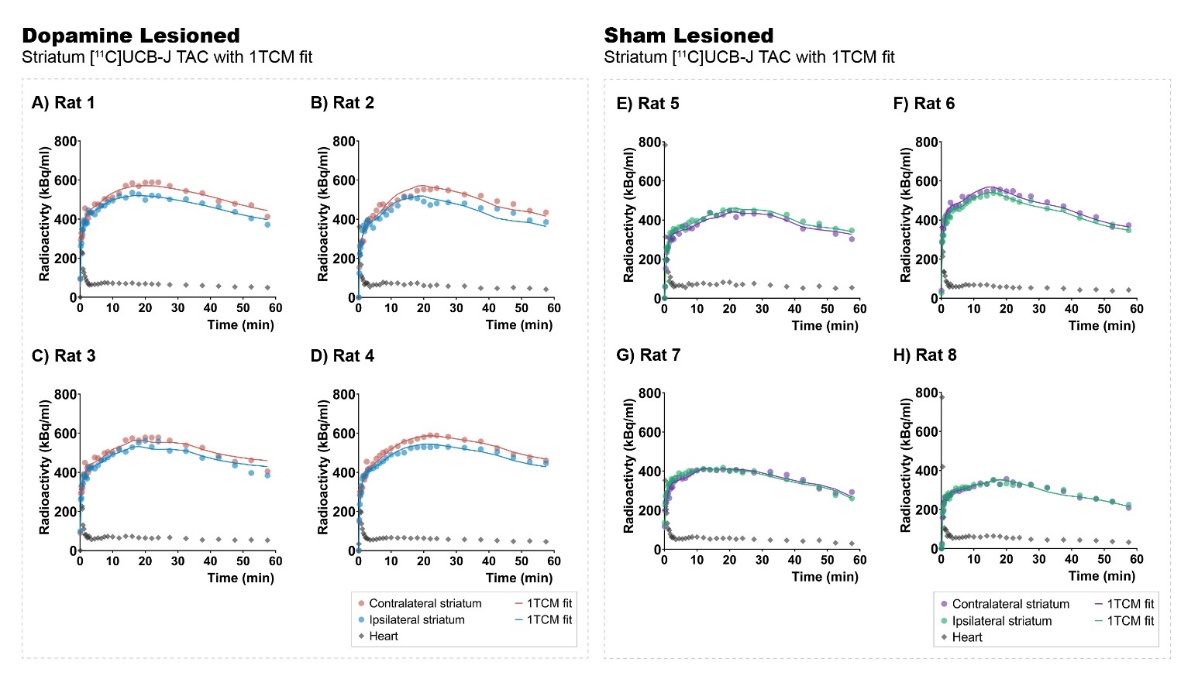

**Supplementary Figure 5.** Representative [^11^C]UCB-J 1TCM model fit in animals within the dopamine and sham lesioned groups. Time activity curves (TAC) demonstrated are from the ipsilateral and contralateral striatum and the heart, used as image-derived input function (IDIF). 1TCM fit using the IDIF is used to extract V_T_ values and other modelling parameters like K1 and k2 values.

#### 2.2 Supplementary Table

**Supplementary Table 1:** Summarised table of [^11^C]UCB-J 1TCM modelling parameters in the ipsilateral and contralateral side of dopamine and sham lesioned animals. Mean values with standard deviation are presented for all animals (n = 4/group). Stri= striatum, DMS= dorsomedial striatum, DLS= dorsolateral striatum, NAc= nucleus accumbens, vMB= ventral midbrain, Thal= thalamus, mPFC= medial prefrontal cortex, ACC= anterior cingulate cortex, OFC= orbitofrontal cortex, MoC= motor cortex.

|  | **[^11^C]UCB-J 1TCM modelling parameters** | | | | | | | |
| --- | --- | --- | --- | --- | --- | --- | --- | --- |
|  | Dopamine Lesioned | | | | Sham Lesioned | | | |
|  | *Ipsilateral Side* | | *Contralateral Side* | | *Ipsilateral Side* | | *Contralateral Side* | |
| **Region** | **K1** | **k2** | **K1** | **k2** | **K1** | **k2** | **K1** | **k2** |
| **Stri** | 1.18 (0.06) | 0.14 (0.01) | 1.18 (0.08) | 0.12 (0.01) | 1.16 (0.19) | 0.16 (0.01) | 1.14 (0.25) | 0.15 (0.01) |
| **DMS** | 1.23 (0.15) | 0.13 (0.02) | 1.3 (0.16) | 0.13 (0.02) | 1.21 (0.23) | 0.16 (0.01) | 1.17 (0.26) | 0.16 (0.01) |
| **DLS** | 1.25 (0.1) | 0.13 (0.01) | 1.26 (0.13) | 0.13 (0.01) | 1.18 (0.24) | 0.15 (0.01) | 1.16 (0.18) | 0.15 (0.01) |
| **NAc** | 1.04 (0.01) | 0.13 (0.01) | 1.05 (0.03) | 0.12 (0.01) | 1.02 (0.07) | 0.16 (0.02) | 0.96 (0.09) | 0.16 (0.03) |
| **vMB** | 1.45 (0.13) | 0.2 (0.02) | 1.5 (0.15) | 0.19 (0.02) | 1.3 (0.22) | 0.21 (0.04) | 1.29 (0.15) | 0.21 (0.02) |
| **Thal** | 1.55 (0.11) | 0.15 (0.01) | 1.5 (0.13) | 0.17 (0.01) | 1.42 (0.26) | 0.17 (0.01) | 1.37 (0.16) | 0.17 (0.01) |
| **mPFC** | 1.21 (0.09) | 0.12 (0.01) | 1.27 (0.1) | 0.13  (0.01) | 1.22 (0.18) | 0.16 (0.01) | 1.23 (0.21) | 0.16 (0.01) |
| **ACC** | 1.06 (0.09) | 0.12 (0.01) | 1.09 (0.09) | 0.13 (0.01) | 0.9 (0.19) | 0.17 (0.01) | 0.78 (0.09) | 0.16 (0.01) |
| **OFC** | 1.06 (0.03) | 0.14 (0.01) | 1.09 (0.06) | 0.15 (0.007) | 1.07 (0.2) | 0.17 (0.01) | 0.99 (0.13) | 0.17 (0.01) |
| **MoC** | 0.98 (0.06) | 0.13 (0.01) | 0.94 (0.08) | 0.13 (0.01) | 1.03 (0.19) | 0.16 (0.01) | 0.96 (0.13) | 0.16 (0.01) |
